# Hyaluronic acid dynamics are regulated by neural crest cells and NEDD4 in the developing cardiac outflow tract valves

**DOI:** 10.64898/2026.11.08.764443

**Authors:** Jasmine Hartmann, Deepti Domingo, Sarah Boyle, Ǫuenten Schwarz, Sophie Wiszniak

**Author notes:** Corresponding author: Sophie Wiszniak.

## Abstract

The extracellular matrix (ECM) is essential for cardiac valve development, with hyaluronic acid (HA) playing a critical role in cardiac cushion formation. While HA is required for atrioventricular valve development, its role in semilunar valve formation remains poorly understood. The ubiquitin ligase NEDD4 is essential for cardiac outflow tract development, particularly within cardiac neural crest cells, however its role in semilunar valve development has been unexplored.

Here, we demonstrate that neural crest-specific deletion of NEDD4 in mice disrupts outflow tract valve development, resulting in abnormal leaflet number and morphology. Analysis of valve ECM revealed a specific reduction in HA within the neural crest-derived valve mesenchyme, while other ECM components such as fibronectin and versican remained unaffected. In contrast, deletion of NEDD4 in endocardium-derived valve mesenchyme did not alter HA abundance. Using NEDD4-deficient neural crest cells, we further demonstrate a cell-autonomous requirement for NEDD4 in regulating HA abundance, with *in vitro* evidence supporting impaired HA synthesis rather than increased turnover of HA in the ECM.

Our findings identify a previously unrecognised role for NEDD4 in regulating HA synthesis in cardiac neural crest cells, revealing a cell-type-specific mechanism governing semilunar valve development and highlighting distinct cellular requirements for ECM regulation across cardiac valve populations.

## Introduction

The extracellular matrix (ECM) is critically important in embryonic development, with dynamic regulation of matrix components controlling diverse aspects of tissue growth, structure, morphogenesis, and remodelling. In the heart, ECM regulation is essential for development of contractile myocardium, formation of the ventricular trabeculae, and the cardiac valves including the atrio-ventricular valves and the semilunar valves of the outflow tract (Del Monte-Nieto et al., 2020). During the initial stages of valve development, acellular ECM known as the cardiac jelly occupies the space between the inner endocardial and outer myocardial layers of the primitive heart tube. As valve development proceeds, the cardiac jelly is invaded by interstitial cells, forming localised thickened swellings known as cardiac cushions that will remodel to form the mature valves. In the atrio-ventricular valves, the cardiac cushions are composed of cells derived from the endocardium that have undergone an endocardial-to-mesenchymal transition (EndMT) to invade the cardiac jelly. In the semilunar valves, in addition to EndMT derived cells, the valve mesenchyme also contains cells of neural crest cell origin (O’Donnell and Yutzey, 2020).

Hyaluronic acid (HA) is one of the most abundant ECM components in the developing cardiac valves. Loss of HA either by excessive degradation (Nakamura and Manasek, 1981) or inhibition of synthesis in *Has2^-/-^* embryos (Camenisch et al., 2000) leads to cardiac valve defects. This is driven in part by loss of ErbB2/ErbB3 signalling activation in valve endocardium and mesenchyme, resulting in disrupted EndMT and cushion formation (Camenisch et al., 2002). While these studies have determined an essential role for HA in atrio-ventricular valve development, early embryonic lethality in these models has precluded investigation of the roles for HA in semilunar valve development.

Prior work has determined an essential role for the ubiquitin ligase NEDD4 in cardiac outflow tract development, with important tissue-restricted roles specifically in the cardiac neural crest (Wiszniak et al., 2026). While NEDD4 has been shown to regulate the paracrine signalling activity of neural crest cells in early stages of cardiac outflow tract lengthening (Wiszniak et al., 2026), the roles for NEDD4 and neural crest cells in semilunar valve development have remained unexplored in this model. Here, we demonstrate dysmorphic outflow tract valve development in *Wnt1-Cre; Nedd4^ff/ff^* embryos, including disrupted leaflet numbers and morphology. Analysis of ECM components in the valve mesenchyme showed a specific reduction in HA, while other ECM components such as fibronectin and versican were unaffected. The reduction in HA was also specific to neural crest-derived valve mesenchyme, as HA dynamics were unaffected in *Mef2cAHF-Cre; Nedd4^ff/ff^* embryos with loss of NEDD4 in the semilunar valve endocardial-derived (EndMT) mesenchyme. Using *Nedd4^-/-^* neural crest cells, we demonstrate cell-autonomous roles for neural crest cells and NEDD4 in regulating HA abundance, with *in vitro* evidence supporting a role for NEDD4 in regulating HA synthesis, rather than turnover. Our findings suggest important cell-type-specific roles for NEDD4 and HA in regulating outflow tract valve development, revealing the critical requirement of cardiac neural crest cells for HA synthesis, and further highlighting the distinct cellular processes that underpin semilunar versus atrioventricular valve development.

## Results and Discussion

### *Wnt1-Cre; Nedd4^ff/ff^* embryos have dysmorphic outflow tract valves

Our prior work discovered an essential role for NEDD4 in cardiac outflow tract development, by demonstrating that loss of *Nedd4* in neural crest cells caused artery-ventricle alignment defects (such as double-outlet right ventricle) due to defective growth of the second heart field-derived outflow tract myocardium (Wiszniak et al., 2026). Additionally, we discovered *Wnt1-Cre; Nedd4^ff/ff^* embryos exhibited hypoplastic semilunar valves and dysmorphic valve leaflets (Wiszniak et al., 2026). We therefore sought to investigate the detailed molecular mechanisms of defective semilunar valve development in *Wnt1-Cre; Nedd4^ff/ff^* embryos.

While the outflow tract cushions develop normally at E11.5, with no significant differences in cushion size in *Wnt1-Cre; Nedd4^ff/ff^* embryos (**Fig. 1A,B**), by E12.5 the outflow tract valves are significantly hypoplastic, disproportionately affecting the pulmonary valve (**Fig. 1C, D**). Dysmorphic valve leaflet numbers are also observed in both the aortic and pulmonary valves (**Fig. 1C, F**), often presenting with only 2 or up to 5 valve leaflets rather than the usual 3 leaflets observed in wildtype embryos. Valve leaflet morphology was also visualised through whole mount imaging of the dissected outflow tract, followed by 3D surface generation in Imaris, which revealed dysmorphous valve leaflets in *Wnt1-Cre; Nedd4^ff/ff^* embryos compared to wildtype at E12.5 (**Fig. 1E**). Lineage tracing of neural crest cells and their derivatives with EGFP using the *Z/EG* reporter demonstrated that *Wnt1-Cre; Z/EG; Nedd4^ff/ff^* embryos do not exhibit any deficiency of neural crest cells in the developing outflow tract from E11.5-E12.5 (**Supp. Fig. 1A-B**), indeed there was a small but significant increase in neural crest cells in the outflow tract mesenchyme at E11.5 in *Wnt1-Cre; Z/EG; Nedd4^ff/ff^* embryos. There were also no changes in cell death or proliferation between genotypes (**Supp. Fig. 1C-F**) that may explain the reduced valve size in *Wnt1-Cre; Nedd4^ff/ff^* embryos. Hence, we propose that NEDD4 plays a critical functional role in neural crest cells once they have populated the outflow tract to direct correct valve morphogenesis.

**Figure 1:**
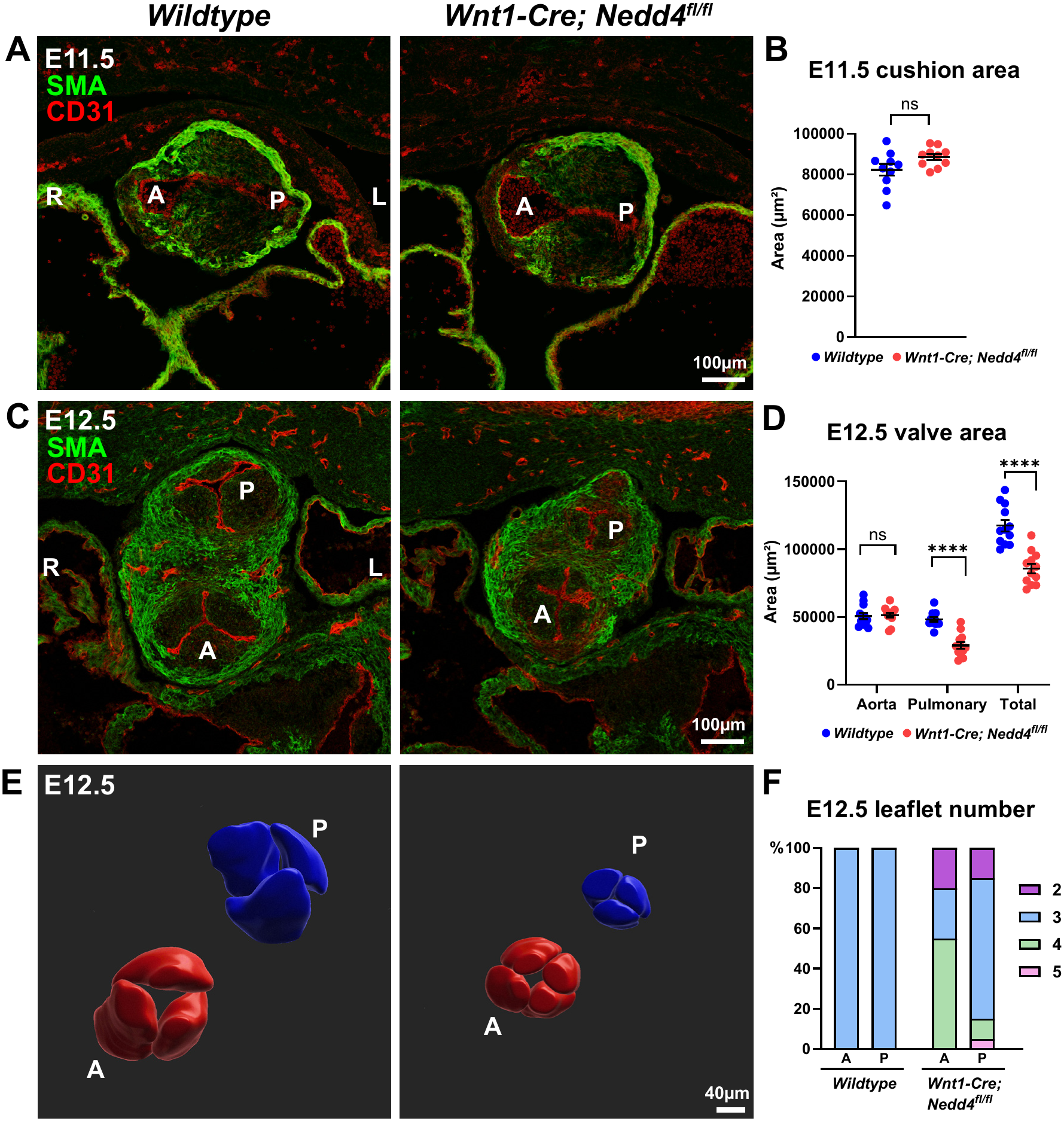
Outflow tract valve development is disrupted in *Wnt1-Cre; Nedd4^ff/ff^* embryos. **A:** Coronal sections through the outflow tract of wildtype and *Wnt1-Cre; Nedd4^ff/ff^* embryos at E11.5, immunostained for smooth muscle actin (SMA) to label vascular smooth muscle and CD31 to label endothelial/endocardial cells. **B:** Outflow tract cushion area at E11.5, measured from 2D sections as in (A). ns, not significant; unpaired t-test. **C:** Coronal sections through the outflow tract at E12.5, immunostained for SMA and CD31. **D:** Outflow tract valve area at E12.5, measured from 2D sections as in (C). ns, not significant; **** p<0.0001; 2 way ANOVA multiple comparisons. **E:** 3D model of outflow tract valves from wildtype and *Wnt1-Cre; Nedd4^ff/ff^* embryos at E12.5, generated through wholemount imaging of the outflow tract, followed by segmentation and surface visualisation using Imaris software. **F:** Number of valve leaflets in aortic and pulmonary valves of wildtype and *Wnt1-Cre; Nedd4^ff/ff^* embryos at E12.5, quantified from sections or 3D models (as in C and E). Graph represents the proportion of total embryos analysed, wildtype n=20, *Wnt1-Cre; Nedd4^ff/ff^* n=20. R, right; L, left; A, aortic; P, pulmonary.

### ECM components are altered in *Wnt1-Cre; Nedd4^ff/ff^* outflow tract valves

We previously performed mRNA sequencing on laser-capture micro-dissected cardiac neural crest and second heart field tissue from E9.5 wildtype and *Wnt1-Cre; Nedd4^ff/ff^* embryos (Wiszniak et al., 2026). Gene ontology analysis of differentially expressed genes identified ‘Extracellular matrix’ as one of the top GO terms (Wiszniak et al., 2026), suggesting ECM components may be disrupted in *Wnt1-Cre; Nedd4^ff/ff^* embryos, and led us to hypothesise that ECM disturbances may underpin the cardiac valve defects observed later in outflow tract development at E11.5-E12.5. The ECM is critical for cardiac development, particularly in the cardiac valves where the proper structural organisation of the cardiac cushions is essential for valve and leaflet maturation (Del Monte-Nieto et al., 2020, Lockhart et al., 2011). HA is one of the most abundant ECM components in the cardiac valves, with critical roles in directing EndMT (Toole, 2001, Camenisch et al., 2000, Camenisch et al., 2002). Fibronectin is also critically important for cushion EndMT (Icardo et al., 1992, Astrof et al., 2007) as well as valve leaflet remodelling and maturation in response to blood flow (Steed et al., 2016). Versican is a proteoglycan that can bind to HA, and is cleaved by proteases into the bioactive NeoVersican fragment which is important for cardiac cushion remodelling (Kern et al., 2006).

Analysis of ECM components in the developing outflow tract cushions and valves at E11.5 and E12.5 revealed that while Fibronectin and NeoVersican remained unchanged (**Fig. 2B, C, E, F**), HA abundance was significantly reduced in *Wnt1-Cre; Nedd4^ff/ff^* embryos at both timepoints (**Fig. 2A, D**). The loss of HA in *Wnt1-Cre; Nedd4^ff/ff^* embryos was specific to the developing semilunar valves, as other cardiac structures that do not contain neural crest cells, such as the atrio-ventricular canal (AVC) and ventricular trabeculae, were unaffected (**Fig. 2A** and **Supp. Fig. 2**). This suggests that neural crest cells, and neural crest-derived NEDD4, play a particularly important role in specifically regulating HA abundance in the developing semilunar valves.

**Figure 2:**
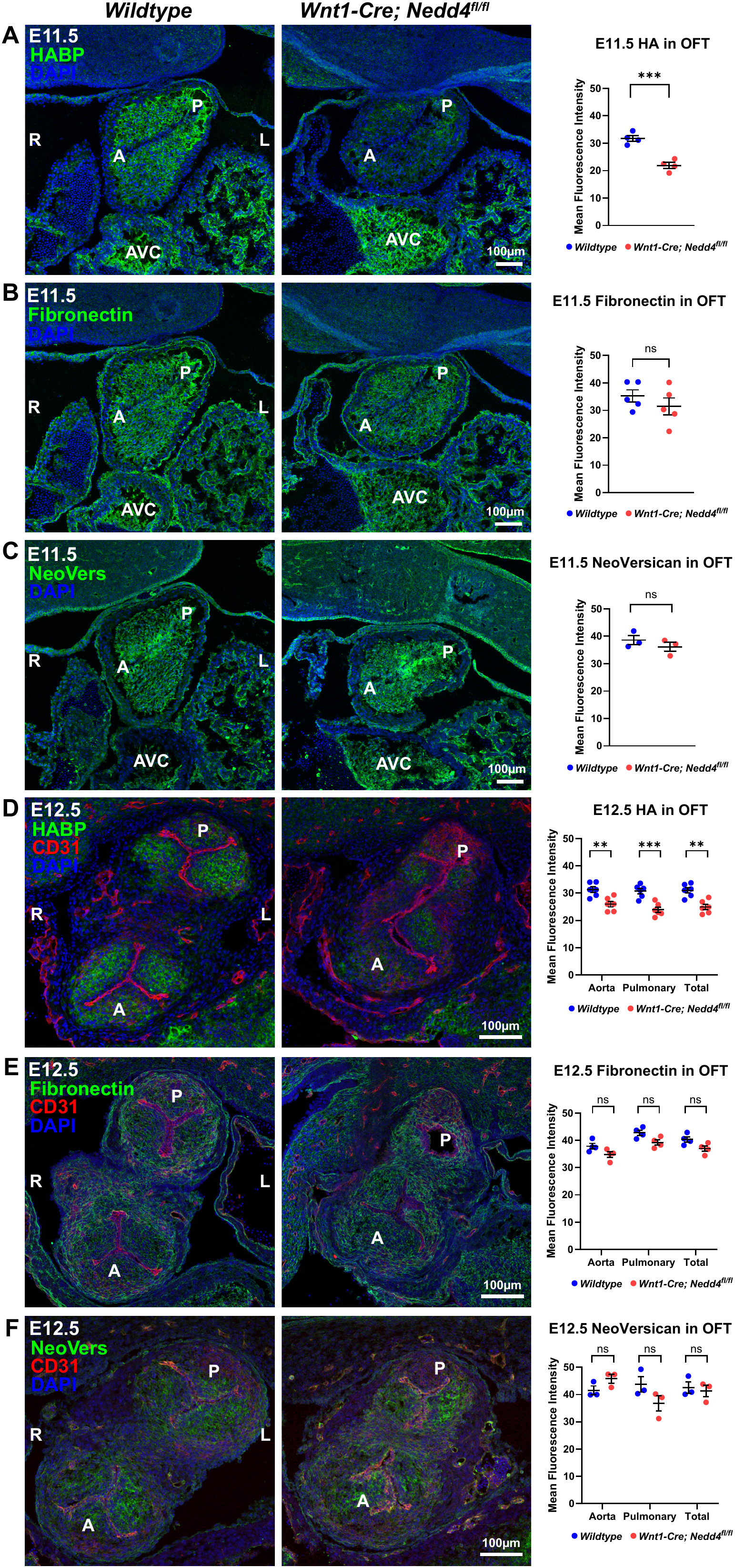
ECM components are altered in *Wnt1-Cre; Nedd4^ff/ff^* outflow tract valves. **A:** Coronal sections through the outflow tract of wildtype and *Wnt1-Cre; Nedd4^ff/ff^* embryos at E11.5, stained with biotinylated hyaluronic acid binding protein (HABP) to label areas of HA. Graph represents quantification of mean fluorescence intensity (MFI) of HA in the outflow tract cushions. *** p<0.001; unpaired t-test. **B:** E11.5 outflow tract sections immunostained for fibronectin, and MFI quantified in graph. ns, not significant; unpaired t-test. **C:** E11.5 outflow tract sections immunostained for neo-versican, and MFI quantified in graph. ns, not significant; unpaired t-test. **D:** E12.5 outflow tract sections immunostained for HABP and CD31, and HA MFI quantified in graph. ** p<0.01; *** p<0.001; 2 way ANOVA multiple comparisons. **E:** E12.5 outflow tract sections immunostained for fibronectin and CD31, and fibronectin MFI quantified in graph. ns, not significant; 2 way ANOVA multiple comparisons. **F:** E12.5 outflow tract sections immunostained for neo-versican and CD31, and neo-versican MFI quantified in graph. ns, not significant; 2 way ANOVA multiple comparisons. R, right; L, left; A, aortic; P, pulmonary; AVC, atrioventricular canal.

### Valve mesenchyme of neural crest origin, but not endocardial origin, is essential for NEDD4-mediated HA production

The semilunar valves of the outflow tract undergo developmental processes that are distinct from the atrioventricular valves, particularly in regard to the developmental origins of mesenchymal cells that populate the cardiac cushions and later remodel into the valve leaflets. In the atrioventricular valves, cushion mesenchyme is derived from EndMT of the overlying endocardium, while in the semilunar valves, the mesenchyme is populated both by cells derived from EndMT, as well as from neural crest origin (O’Donnell and Yutzey, 2020). The *Mef2cAHF-Cre* driver is widely used to lineage trace cells of second heart field origin, but is also expressed in the endocardium of the semilunar valves (Verzi et al., 2005), and hence is amenable to lineage tracing cells of EndMT origin in the valve cushions. At E11.5, the semilunar valve mesenchyme is predominantly neural crest-derived, as demonstrated by lineage tracing of neural crest-derived tissue with *Wnt1-Cre; Z/EG*, with very minor contributions from the *Mef2cAHF-Cre; Z/EG* lineage-traced endocardium (**Fig. 3A**). However by E12.5, EndMT-derived cells have now begun to invade the valve mesenchyme with valve leaflets comprised of lineage-traced cells of both neural crest and endocardial origin (**Fig. 3B**).

**Figure 3.**
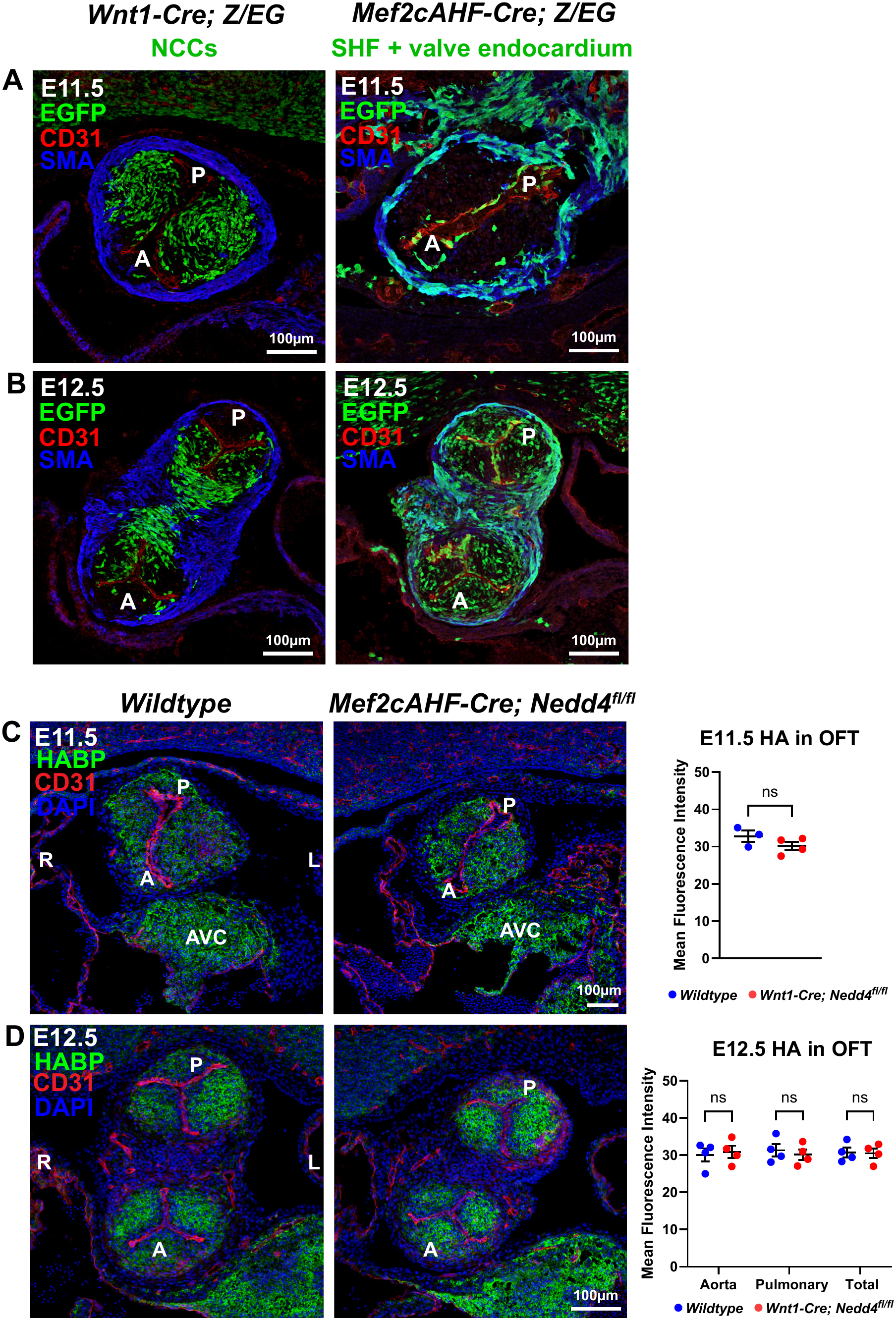
HA abundance is unaffected in EndMT-derived valve mesenchyme upon loss of NEDD4. **A:** Coronal sections through the outflow tract of *Wnt1-Cre; Z/EG* and *Mef2cAHF-Cre; Z/EG* embryos at E11.5, immunostained for EGFP to label lineage-traced cells, CD31 and smooth muscle actin (SMA). At E11.5, the outflow tract valve mesenchyme is predominantly neural crest-derived, with minimal contribution of endocardial-derived cells. **B:** Sections through the outflow tract of E12.5 *Wnt1-Cre; Z/EG* and *Mef2cAHF-Cre; Z/EG* embryos, immunostained for EGFP, CD31 and SMA. At E12.5, the outflow tract valve mesenchyme is populated by cells of both neural crest and endocardial origin. **C:** Sections through the outflow tract of wildtype and *Mef2cAHF-Cre; Nedd4^ff/ff^* embryos at E11.5, stained with biotinylated hyaluronic acid binding protein (HABP) to label areas of HA, and CD31. Graph represents quantification of mean fluorescence intensity (MFI) of HA in the outflow tract cushions. ns, not significant; unpaired t-test. **D:** E12.5 wildtype and *Mef2cAHF-Cre; Nedd4^ff/ff^* outflow tract sections stained with HABP and CD31, and HA MFI quantified in graph. ns, not significant; 2 way ANOVA multiple comparisons. R, right; L, left; A, aortic; P, pulmonary; AVC, atrioventricular canal.

We assessed HA abundance in the outflow tract of *Mef2cAHF-Cre; Nedd4^ff/ff^* embryos to determine if NEDD4 is required in EndMT-derived mesenchyme, in addition to neural crest-derived mesenchyme, for production of HA. Unlike in *Wnt1-Cre; Nedd4^ff/ff^* embryos, no significant differences were observed in HA abundance in the outflow tract of *Mef2cAHF-Cre; Nedd4^ff/ff^* embryos when compared to wildtype controls, at both E11.5 and E12.5 (**Fig. 3C,D**). Furthermore, valve morphology appeared normal in *Mef2cAHF-Cre; Nedd4^ff/ff^* embryos, which is consistent with our prior observations of a lack of cardiac outflow tract defects in this model (Wiszniak et al., 2026). Together, our findings suggest that the predominant role for NEDD4 in regulating abundance of HA in the developing outflow tract valves occurs in cells of neural crest-origin, and highlights the critical role that neural crest cells play in regulation of ECM dynamics in the semilunar valves, particularly during the early stages of valve development prior to endocardial invasion of the valve mesenchyme.

Previous studies focussed on atrioventricular valve development have shown that HA functions to activate ErbB2/ErbB3 signalling to facilitate correct endocardial cushion formation (Camenisch et al., 2002). Consistent with this, we observed a significant reduction in ErbB2 in the outflow tract cushions of *Wnt1-Cre; Nedd4^ff/ff^* embryos at E11.5 (**Supp Fig. 3A**), yet ErbB2 levels were only marginally and not significantly reduced at E12.5 (**Supp Fig. 3B**), and no significant differences were observed in EndMT markers including SLUG and *Snai1* at E11.5 (**Supp Fig. 4**). Hence our data suggest that reduced HA abundance in the outflow tract cushions of *Wnt1-Cre; Nedd4^ff/ff^* embryos has a minor effect on ErbB2 signalling, yet does not affect EndMT, and that a reduction in EndMT does not explain the outflow tract valve hypoplasia in this model. This may indicate important differences in EndMT regulatory mechanisms between the atrioventricular and outflow tract valves, as well as reflecting the differences in phenotypes observed between decreased HA abundance in *Wnt1-Cre; Nedd4^ff/ff^* embryos versus complete loss of HA in *Has2^-/-^* models (Camenisch et al., 2002, Camenisch et al., 2000).

### Neural crest cells regulate HA dynamics in a cell-autonomous manner

NEDD4 is abundantly expressed in the neural crest-derived outflow tract cushions and valves at E11.5-E12.5 (**Supp Fig. 5**), raising the possibility that neural crest cells are a major source of HA, and that NEDD4 may regulate HA dynamics during outflow tract valve development. To investigate the cell-autonomous roles for NEDD4 and neural crest cells in HA synthesis and degradation, we used previously generated *Nedd4^+/+^* and *Nedd4^-/-^* NCU10K neural crest cell lines (Lohraseb et al., 2022) to investigate ECM dynamics.

To analyse cellular production of ECM, NCU10K cells were grown for 6 days in the presence of ascorbic acid to promote ECM synthesis and maturation (Kaukonen et al., 2017). Cells were then either fixed intact, or the cells were first removed from the underlying matrix to leave de-cellularised ECM (dECM) for analysis (**Fig. 4A**). NCU10K parental cells produce high levels of HA, both in intact cells, as well as in the deposited dECM (**Fig. 4B**). While two independent NCU10K *Nedd4^+/+^* control CRISPR cell lines produce HA at levels indistinguishable from the parental line, two *Nedd4^-/-^* cell lines demonstrated significantly reduced HA production, evident through imaging of both intact cells and in dECM (**Fig. 4B, E**). NCU10K cells produced limited amounts of Fibronectin and NeoVersican, and consistent with the *in vivo* observations in *Wnt1-Cre; Nedd4^ff/ff^* embryos, no differences in Fibronectin or NeoVersican abundance were observed between *Nedd4^+/+^* and *Nedd4^-/-^* cells (**Supp Fig. 6**). This suggests that NEDD4 plays a specific and important role in regulating the abundance of HA in neural crest cells in a cell-autonomous manner.

**Figure 4:**
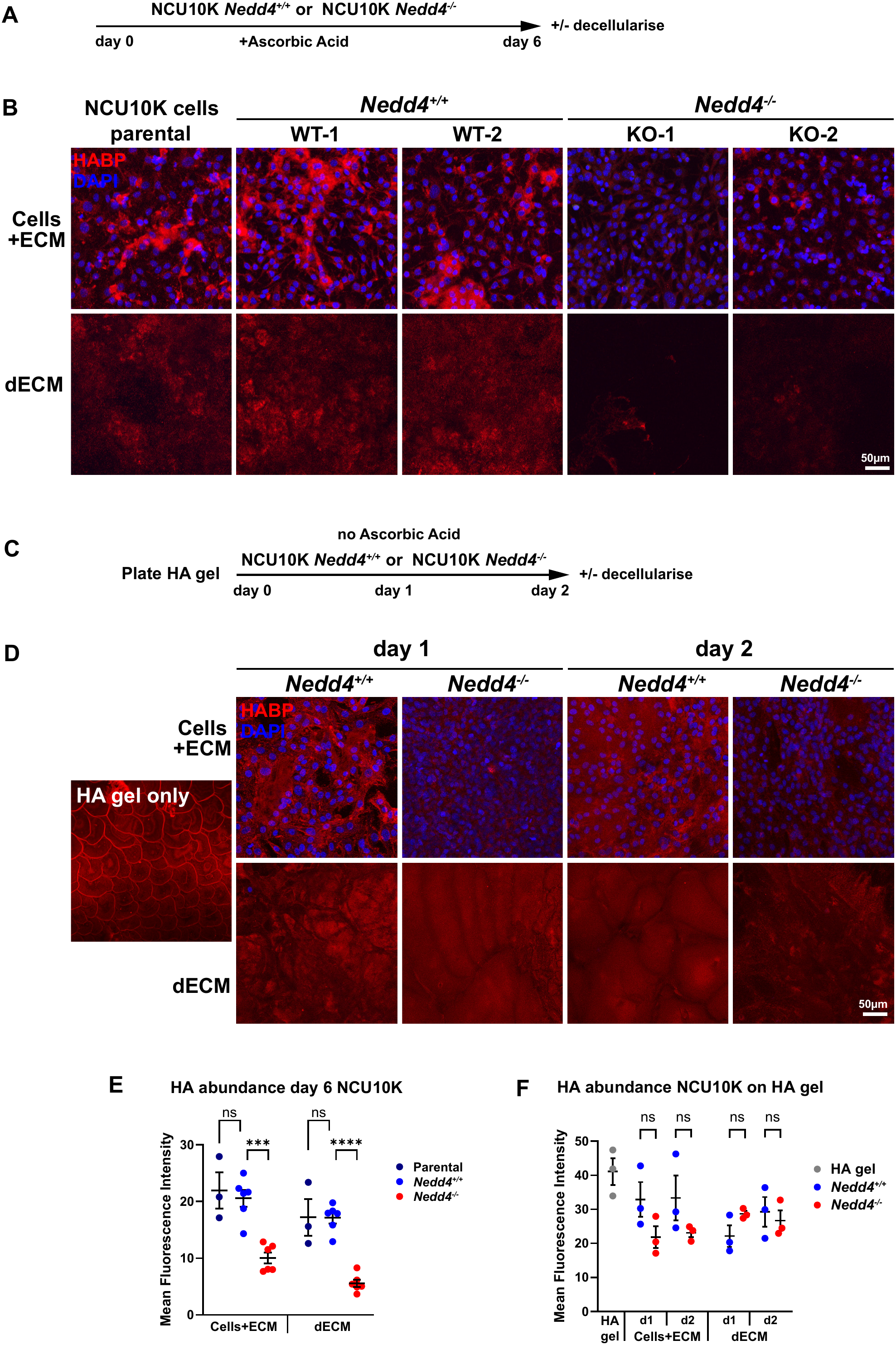
NEDD4 cell-autonomously regulates HA synthesis in neural crest cells. **A:** Diagram of experimental procedure, culture conditions and timing of cell fixation/decellularisation. NCU10K parental or CRISPR *Nedd4^+/+^* (WT-1 or WT-2) or CRISPR *Nedd4^-/-^* (KO-1 or KO-2) were cultured for 6 days with ascorbic acid, then fixed before or after decellularisation. **B:** Staining of fixed cells or decellularised ECM (dECM) with biotinylated hyaluronic acid binding protein (HABP) to label areas of HA. Panels include NCU10K parental cells, two *Nedd4^+/+^* (WT-1 or WT-2) CRISPR control lines, and two *Nedd4^-/-^* (KO-1 or KO-2) CRISPR knockout lines. **C:** Diagram of experimental procedure. Recombinant HA gel was first plated in culture dishes, and *Nedd4^+/+^* or *Nedd4^-/-^* cells were cultured for 1 or 2 days without ascorbic acid, then fixed before or after decellularisation. **D:** Staining of HA gel only, or fixed cells or dECM with HABP. **E:** Graph of HA mean fluorescence intensity (MFI), quantified from images as in (B). Data from the two *Nedd4^+/+^* (WT-1 and WT-2) and two *Nedd4^-/-^* (KO-1 or KO-2) cell lines were combined to give 6 biological replicates in total. ns, not significant; *** p<0.001; **** p<0.0001; 2 way ANOVA multiple comparisons. **F:** Graph of HA mean fluorescence intensity (MFI), quantified from images as in (D). ns, not significant; 2 way ANOVA multiple comparisons.

We hypothesised the reduced HA abundance observed in *Nedd4^-/-^* cells and cell-derived dECM could be due to reduced synthesis of HA, and/or enhanced degradation of deposited HA in the ECM. To investigate modification of existing ECM by *Nedd4^+/+^* and *Nedd4^-/-^* cells, we first generated cell-derived dECM by culturing either the parental NCU10K cells, or alternatively the ATDC5 chondrogenic cell line which we previously determined to generate HA ECM, for 6 days in the presence of ascorbic acid, followed by decellularisation. *Nedd4^+/+^* or *Nedd4^-/-^* cells were then plated on top of this dECM, and cultured for only 2 days in the absence of ascorbic acid to limit cell-autonomous HA synthesis. Cells were then either fixed intact, or removed to leave dECM to assess whether *Nedd4^+/+^* and *Nedd4^-/-^* cells exhibited differences in their ability to degrade existing ECM (**Supp. Fig. 7A,D**). While we did not observe significant differences in HA abundance in dECM on which *Nedd4^+/+^* and *Nedd4^-/-^* cells were cultured (**Supp. Fig. 7B,C,E,F**), the extensive washing steps required to prepare the unfixed NCU10K or ATDC5 dECM for subsequent culture of live cells resulted in limited remaining dECM for analysis. Hence, rather than cell-derived dECM, we used commercially available recombinant HA hydrogels to directly assess the ability of *Nedd4^+/+^* or *Nedd4^-/-^* cells to modify extracellular HA. HA hydrogels were prepared and plated into culture dishes, with *Nedd4^+/+^* or *Nedd4^-/-^* cells grown on top of the gel for 1 or 2 days, followed by fixing of intact cells or decellularisation to assess potential modification of the exogenous HA gel (**Fig. 4C**). Analysis of HA abundance in the decellularised gel following culture with *Nedd4^+/+^* or *Nedd4^-/-^* cells revealed no significant differences between genotypes, both at 1 and 2 days of culture (**Fig. 4D,F**). This suggests that *Nedd4^+/+^* and *Nedd4^-/-^* cells do not exhibit differences in their ability to turnover HA in existing ECM. Taken together, our data supports the hypothesis that NEDD4 regulates abundance of HA in neural crest-derived tissue in a cell-autonomous manner, through a mechanism affecting the synthesis of HA, rather than through modifying HA turnover in the ECM.

Ubiquitination has been shown to play important roles in HA synthesis, particularly through regulation of the key HA synthesis enzyme HAS2. Catalytic activity of HAS2 is dependent on monoubiquitination at Lys-190 (Karousou et al., 2010). Deubiquitinating enzymes also regulate HAS2 activity, with USP17 removing polyubiquitination and enhancing stability of HAS2 and HA production, while USP4 preferentially removes monoubiquitination and reduces HA synthesis (Mehic et al., 2017). While the molecular mechanisms linking NEDD4 to HA synthesis remain undefined, our work implicates a functional role for NEDD4 in the HA biosynthesis pathway for the first time, expanding knowledge of the role for the ubiquitin pathway in HA regulation.

## Conclusion

Here, we demonstrate a critical role for neural crest cells in regulating the synthesis of HA in the developing outflow tract valves, via a mechanism that is dependent on the function of the ubiquitin ligase NEDD4. Our findings reveal important cell-type-specific roles for NEDD4 and HA in regulating outflow tract valve development, with differential requirements of HA synthesis from neural crest versus endocardial origin. The semilunar valves of the developing outflow tract are unique in their constituents, comprising cells of both neural crest and endocardial origin, distinguishing them from the atrioventricular valves which contain only endocardial-derived cells. Our work therefore demonstrates that divergent cellular processes underpin semilunar and atrioventricular valve development, and highlights the unique contribution and instructive roles that neural crest cells play in the outflow tract valves, which has important impacts for mechanistic understanding of valve development and disease.

## Materials and Methods

### Mice

All experiments were carried out in accordance with ethical guidelines of the Adelaide University Animal Ethics Committee. Mice are housed with a 12h dark/light cycle, with temperature maintained between 19-23°C and 40-70% relative humidity. Mice used for timed matings were between 6-12 weeks of age. All mouse lines were maintained on a mixed C57BL/6 background. *Nedd4^ff/ff^* (Kawabe et al., 2010), *Wnt1-Cre* (Danielian et al., 1998) and *Mef2cAHF-Cre* (Verzi et al., 2005) mouse lines have been described previously. To remove *Nedd4* in specific tissues, we crossed *Nedd4^ff/ff^* females to *Wnt1-Cre;Nedd4^ff/+^* or *Mef2cAHF-Cre;Nedd4^ff/+^* males. The *Z/EG* (Novak et al., 2000) reporter line was also intercrossed with *Wnt1-Cre;Nedd4^ff/+^* and *Mef2cAHF-Cre* mice for lineage tracing. To obtain mouse embryos of defined gestational ages, mice were mated in the evening and the morning of vaginal plug formation was counted as E0.5. In all experiments, stage-matched embryo littermates were selected for comparison using stage-appropriate parameters such as somite number, crown-rump length, and forelimb/digit maturity. For embryonic analyses we did not genotype for sex.

### Immunostaining

Embryos were fixed in 4% paraformaldehyde in PBS. For immunolabelling, cryosections, dissected outflow tract tissue, or fixed cells/dECM samples were blocked in 10% DAKO block, 0.2% BSA, 0.2% Triton X-100 in PBS, and incubated with the indicated primary antibodies. Antibodies used were: mouse anti-alpha smooth muscle actin (Sigma A2547) 1:2000; rat anti-CD31 (Biolegend 102502) 1:150; chicken anti-GFP (Abcam ab13970) 1:1000; rabbit anti-SOX9 (Millipore AB5535) 1:1000; sheep anti-Fibronectin (RCD Systems AF1918) 1:200; rabbit anti-NeoVersican (ThermoFisher PA11748A) 1:1000; rabbit anti-ERBB2 (Cell Signaling Technology 2243) 1:100; rabbit anti-SLUG (Cell Signalimg Technology 9585) 1:100; rabbit anti-NEDD4 (Abcam 14592) 1:300; rabbit anti-phospho-Histone H3 (Millipore 06-570) 1:500; rabbit anti-cleaved-Caspase-3 (Cell Signaling Technology 9661) 1:500; Biotinylated HABP (Merck 385911) 1:200. Slides were mounted in Fluoro-mount G with DAPI (ProSciTech). Confocal images were acquired with a Zeiss LSM 800 microscope and ZEN 2.6 (Blue edition) software.

### Fluorescence intensity and image quantitation

Mean fluorescence intensity (MFI) was measured using ZEN 3.4 (Blue edition) software (Zeiss) and represents the total fluorescence intensity of the region of interest, normalised to the area of the region of interest. For OFT ECM MFI measurements, the entire area of the cushions at E11.5, or the whole leaflets of the semilunar valves at E12.5, were traced and measured. A minimum of 3 and up to 8 sections per embryo were used to calculate MFI as described above, and averaged. Individual points on graphs represent the mean value calculated from 1 embryo. A minimum of 3 and up to 6 embryos per genotype were used for ECM MFI measurements. Additionally, MFI was measured in the AVC, by tracing the entire area of the AVC and measuring MFI as above from a minimum of 3 sections per embryo, and from 3-4 embryos per genotype.

For fixed cells and dECM, HABP MFI was calculated by taking MFI measurements from 3x randomly selected 10,000μm^2^ areas per image, from 3 images per well, from 3x technical replicates (individual wells) per experiment, and averaged to represent an individual point on the relevant graph. Each experiment was performed a minimum of 3 and up to 6 times (biological replicates).

For EGFP, PHH3, cleaved-Caspase-3, SLUG and *Snai1* quantification, the entire area of the OFT cushions (E11.5) or leaflets (E12.5) were traced, and the number of +ve cells counted within the region of interest, and expressed as the number of +ve cells per mm^2^. A minimum of 3 and up to 8 sections per embryo were assessed, and averaged. Individual points on graphs represent the mean value calculated from 1 embryo. A minimum of 3 embryos per genotype were used.

### 3D modelling of OFT

The outflow tract was microdissected from E12.5 embryos and immunostained with SOX9, CD31 and SMA primary antibodies as detailed in ‘Immunostaining’ above, then mounted in low-melt agarose, cleared with benzyl alcohol and benzyl benzoate (BABB), and imaged with a Zeiss LSM 800 microscope and ZEN 2.6 (Blue edition) software. Acquired Z-stack images were imported into Imaris x64 9.7.2 software and 3D rendered surfaces of valve leaflets were drawn by manual segmentation. SOX9 was used as a marker of valve cushion mesenchyme, with leaflets defined by the border of CD31-positive endocardium.

### In situ hybridisation

Cryosections were hybridised with digoxigenin-labelled antisense probes for *Snai1*. Signal was detected using anti-DIG-AP conjugated Fab fragments (Roche), and stained with NBT/BCIP (Roche). Sections were imaged with an Olympus IX73.

### Cell culture

The NCU10K neural crest cell line has been described previously (Wiszniak et al., 2013), and was maintained in DMEM + 10% FBS. Generation of *Nedd4^+/+^* (WT-1 and WT-2) and *Nedd4^-/-^* (KO-1 and KO-2) CRISPR engineered NCU10K cells has been described previously (Lohraseb et al., 2022). To promote production of ECM, NCU10K cells were grown in media supplemented with 50μg/ml ascorbic acid for 6 days. To generate decellularized extracellular matrix (dECM), cells were treated with extraction buffer (0.58% (296mM) ammonium hydroxide, 0.5% Triton X-100 in PBS) following protocols as described (Kaukonen et al., 2017). Whole cells or dECM were fixed in 4% PFA for 20 minutes, then labelled with primary antibodies or HABP as per immunostaining methods described above.

To assess effects of *Nedd4^+/+^* and *Nedd4^-/-^* NCU10K cells on already established dECMs, dECMs were generated from parental NCU10K or ATDC5 chondrogenic cells by culturing for 6 days in ascorbic acid supplemented media, and decellularising following the methods described above. *Nedd4^+/+^* or *Nedd4^-/-^* NCU10K cells were plated directly onto the dECMs, and cultured for 2 days in growth media without ascorbic acid to limit production of ECM by the NCU10K cells. Cells or dECMs were then fixed and labelled as above.

To assess effects of *Nedd4^+/+^* and *Nedd4^-/-^* NCU10K cells on HA gels, HA gel (HyStem-C Scaffold Kit, Sigma-Aldrich, HYSC020) was plated following manufacturers recommendations, and *Nedd4^+/+^* or *Nedd4^-/-^* NCU10K cells were plated directly on top, cultured for 1 or 2 days, then fixed and labelled as above.

### Statistics

In all graphical representations, individual values represent independent biological replicates. Unless otherwise specified, *P* values were calculated using unpaired two-tailed *t* test. For grouped analysis, significance was assessed using 1 or two-way ANOVA multiple comparisons. All graphs and statistical analyses were performed using GraphPad Prism 10.

## Acknowledgements

This research was supported by funds from the National Health and Medical Research Council (NHMRC): Ideas Grant (ID2030340) (S.W. and Ǫ.S.), and Australian Government Research Training Program (RTP) Scholarship (J.H.).

## Figure legends

**Supplementary Figure 1:**
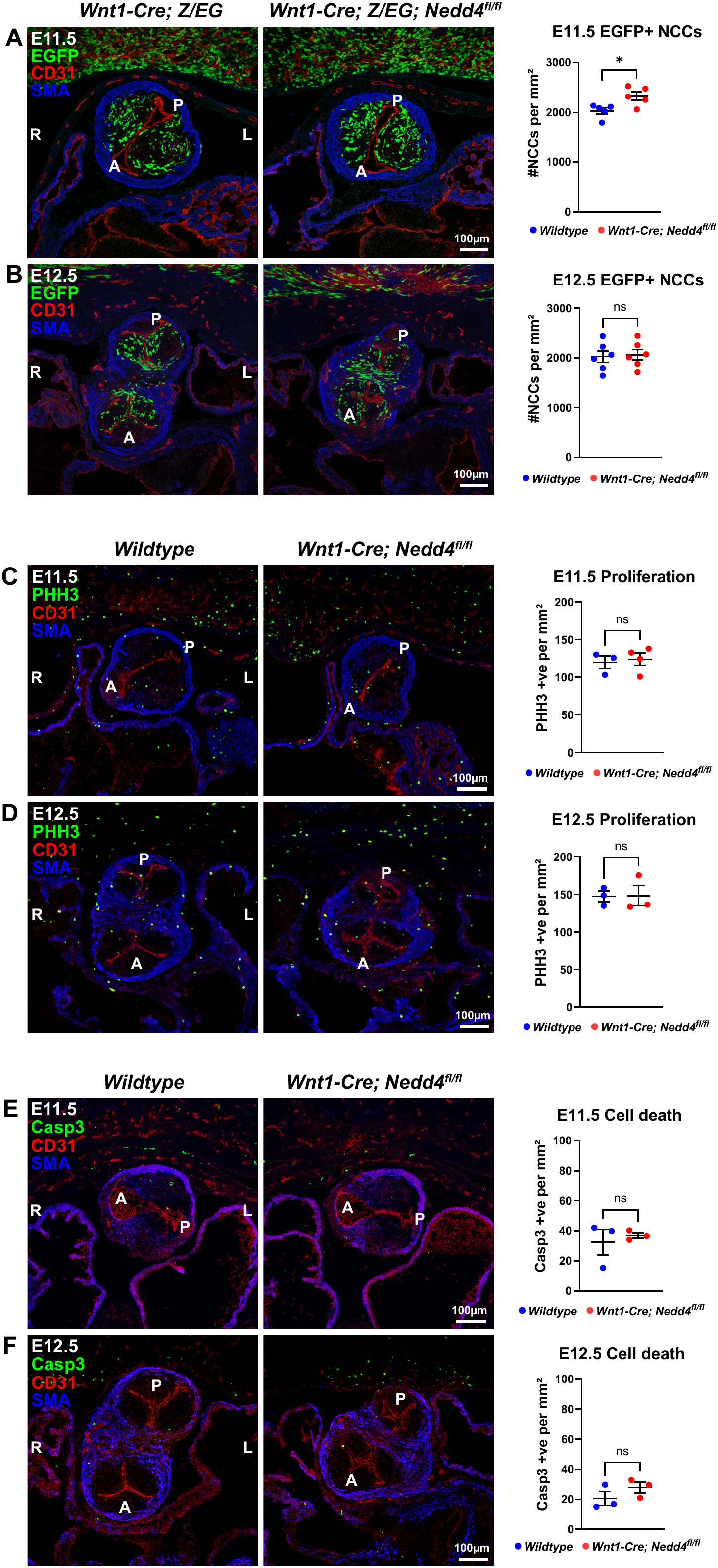
Neural crest cell abundance, proliferation and cell death are unaffected in the outflow tract of *Wnt1-Cre; Nedd4^ff/ff^* embryos. **A:** Coronal sections through the outflow tract of *Wnt1-Cre; Z/EG* and *Wnt1-Cre; Z/EG; Nedd4^ff/ff^* embryos at E11.5, immunostained for EGFP to label lineage-traced neural crest cells, smooth muscle actin (SMA) and CD31. Graph represents quantification of number of EGFP-positive neural crest cells per mm^2^ of outflow tract cushion mesenchyme. * p<0.05; unpaired t-test. **B:** E12.5 sections immunostained for EGFP, CD31 and SMA, and graph representing quantification of EGFP-positive neural crest cells in the valve cushion mesenchyme. ns, not significant; unpaired t-test. **C:** E11.5 sections of wildtype and *Wnt1-Cre; Nedd4^ff/ff^* embryos immunostained for phospho-histone H3 (PHH3), CD31 and SMA, and graph representing quantification of PHH3-positive cells per mm^2^ of outflow tract cushion mesenchyme. ns, not significant; unpaired t-test. **D:** E12.5 sections immunostained for PHH3, CD31 and SMA, and PHH3-positive cells quantified in graph. ns, not significant; unpaired t-test. **E:** E11.5 sections of wildtype and *Wnt1-Cre; Nedd4^ff/ff^* embryos immunostained for cleaved-caspase 3 (Casp3), CD31 and SMA, and graph representing quantification of Casp3-positive cells per mm^2^ of outflow tract cushion mesenchyme. ns, not significant; unpaired t-test. **F:** E12.5 sections immunostained for Casp3, CD31 and SMA, and Casp3-positive cells quantified in graph. ns, not significant; unpaired t-test. R, right; L, left; A, aortic; P, pulmonary.

**Supplementary Figure 2:**
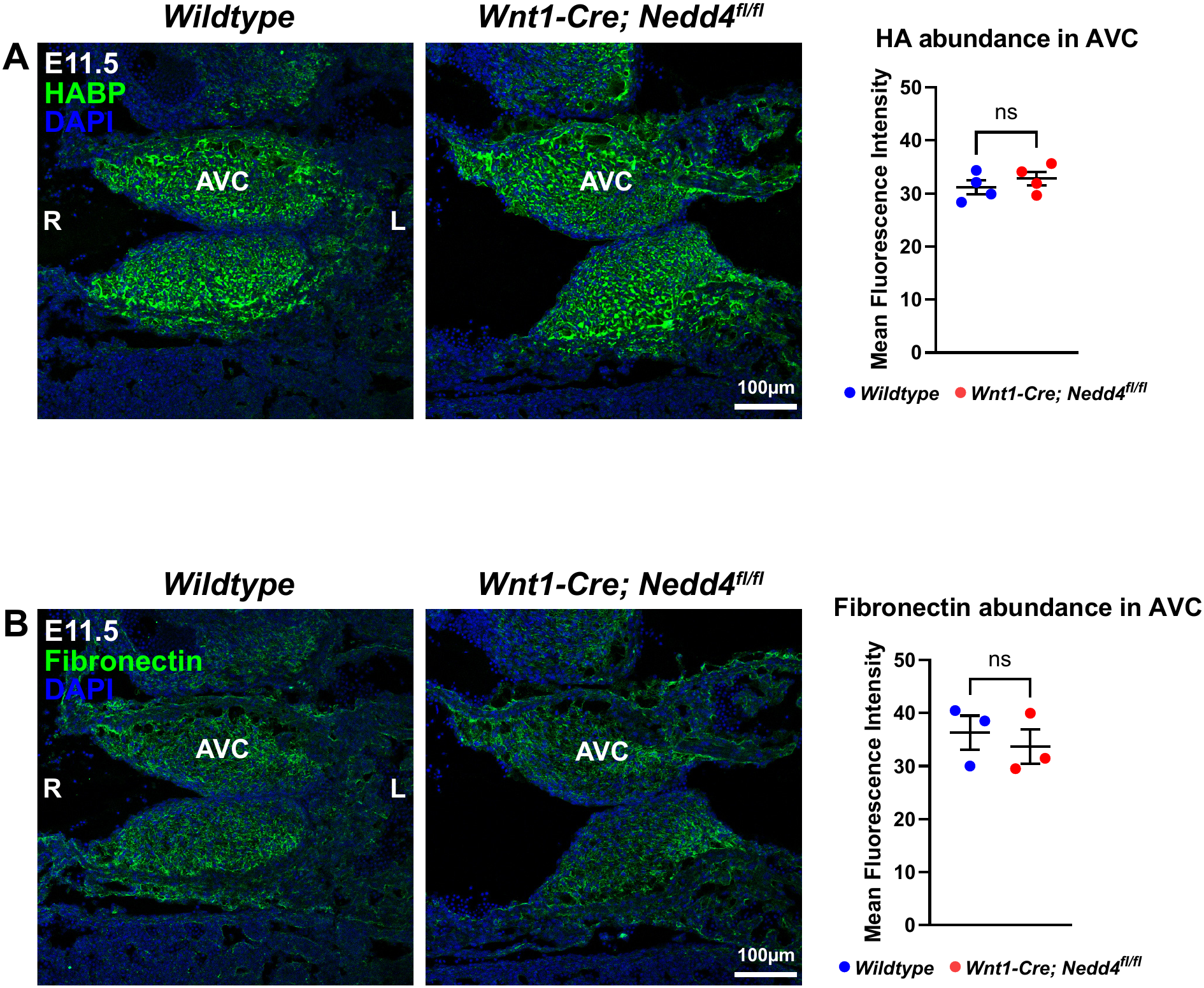
ECM components are unaffected in the atrioventricular canal valve mesenchyme in *Wnt1-Cre; Nedd4^ff/ff^* embryos. **A:** Coronal sections through the atrioventricular (AV) valve of wildtype and *Wnt1-Cre; Nedd4^ff/ff^* embryos at E11.5, stained with biotinylated hyaluronic acid binding protein (HABP) to label areas of HA. Graph represents quantification of mean fluorescence intensity (MFI) of HA in the AV valve mesenchyme. ns, not significant; unpaired t-test. **B:** E11.5 AV valve sections immunostained for fibronectin, and MFI quantified in graph. ns, not significant; unpaired t-test. R, right; L, left; AVC, atrioventricular canal.

**Supplementary Figure 3:**
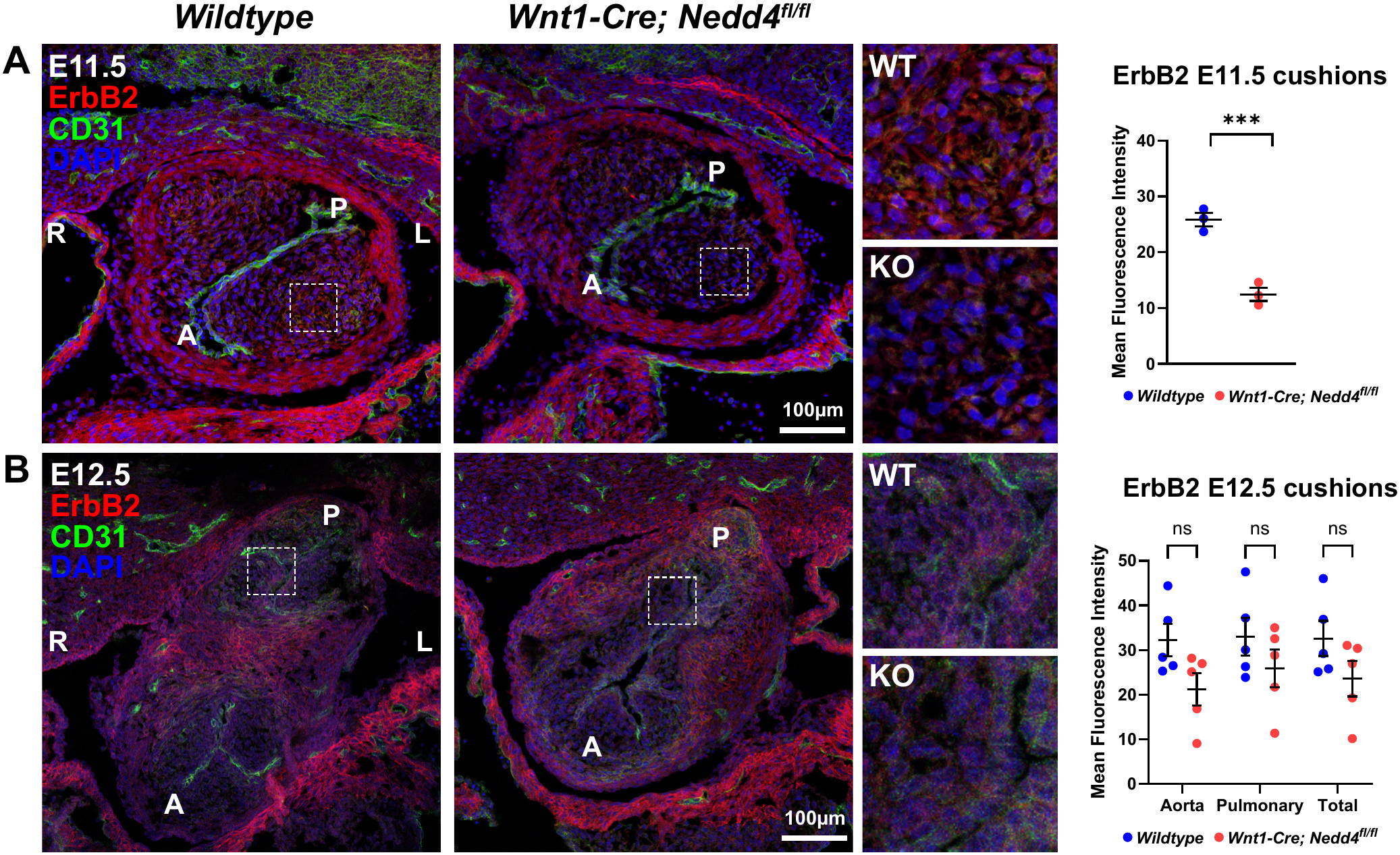
ErbB2 abundance is reduced in the outflow tract mesenchyme of *Wnt1-Cre; Nedd4^ff/ff^* embryos at E11.5. **A:** Coronal sections through the outflow tract of wildtype and *Wnt1-Cre; Nedd4^ff/ff^* embryos at E11.5, immunostained for ErbB2 and CD31. Inset images show higher magnification of the boxed areas. Graph represents quantification of mean fluorescence intensity (MFI) of ErbB2 in the outflow tract cushion mesenchyme. *** p<0.001; unpaired t-test. **B:** E12.5 sections immunostained for ErbB2 and CD31, inset images show higher magnification of the boxed areas. Graph represents MFI of ErbB2 in the outflow tract valves. ns, not significant; unpaired t-test. R, right; L, left; A, aortic; P, pulmonary.

**Supplementary Figure 4:**
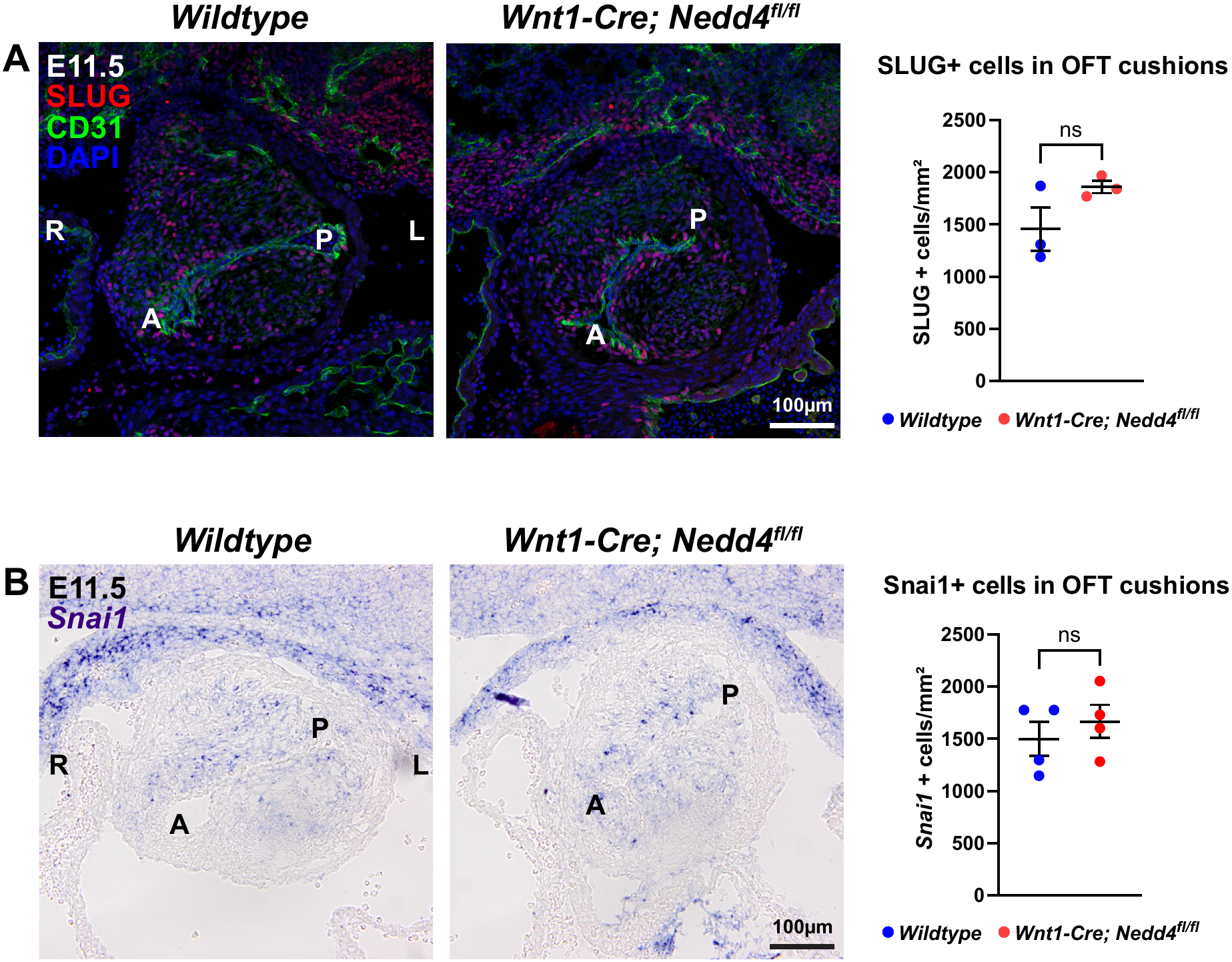
EndMT markers are unchanged in *Wnt1-Cre; Nedd4^ff/ff^* embryos at E11.5. **A:** Coronal sections through the outflow tract of wildtype and *Wnt1-Cre; Nedd4^ff/ff^* embryos at E11.5, immunostained for SLUG and CD31. Graph represents quantification of number of SLUG-positive cells per mm^2^ of outflow tract cushion mesenchyme. ns, not significant; unpaired t-test. **B:** *In situ* hybridisation of E11.5 coronal sections of wildtype and *Wnt1-Cre; Nedd4^ff/ff^* embryos for *Snai1*. Graph represents quantification of number of *Snai1*-positive cells per mm^2^ of outflow tract cushion mesenchyme. ns, not significant; unpaired t-test. R, right; L, left; A, aortic; P, pulmonary.

**Supplementary Figure 5:**
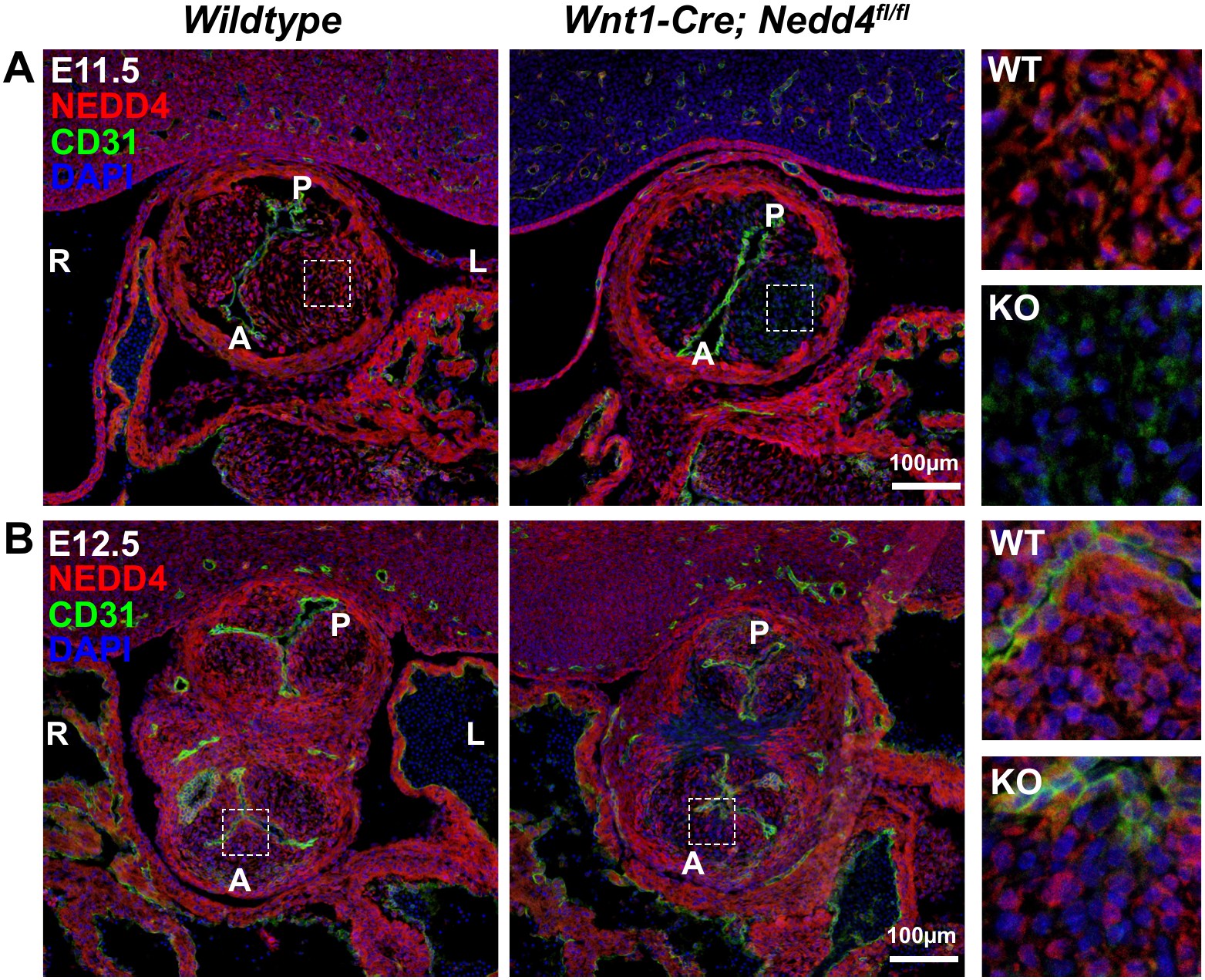
NEDD4 is expressed in the outflow tract including in neural crest-derived valve mesenchyme. **A:** Coronal sections through the outflow tract of wildtype and *Wnt1-Cre; Nedd4^ff/ff^* embryos at E11.5, immunostained for NEDD4 and CD31. Inset images show higher magnification of the boxed areas. NEDD4 is highly expressed in the outflow tract cushion mesenchyme in wildtype, and is mostly absent in the valve mesenchyme in *Wnt1-Cre; Nedd4^ff/ff^* embryos, as expected. **B:** E12.5 sections immunostained for NEDD4 and CD31. Inset images show higher magnification of the boxed areas. NEDD4 is highly expressed throughout the outflow tract and valve leaflets in wildtype, but is absent in some cells of the valve leaflets (the neural crest-derived proportion, while the EndMT-derived proportion still expresses NEDD4), as well as being absent the septum, in *Wnt1-Cre; Nedd4^ff/ff^* embryos. R, right; L, left; A, aortic; P, pulmonary.

**Supplementary Figure 6:**
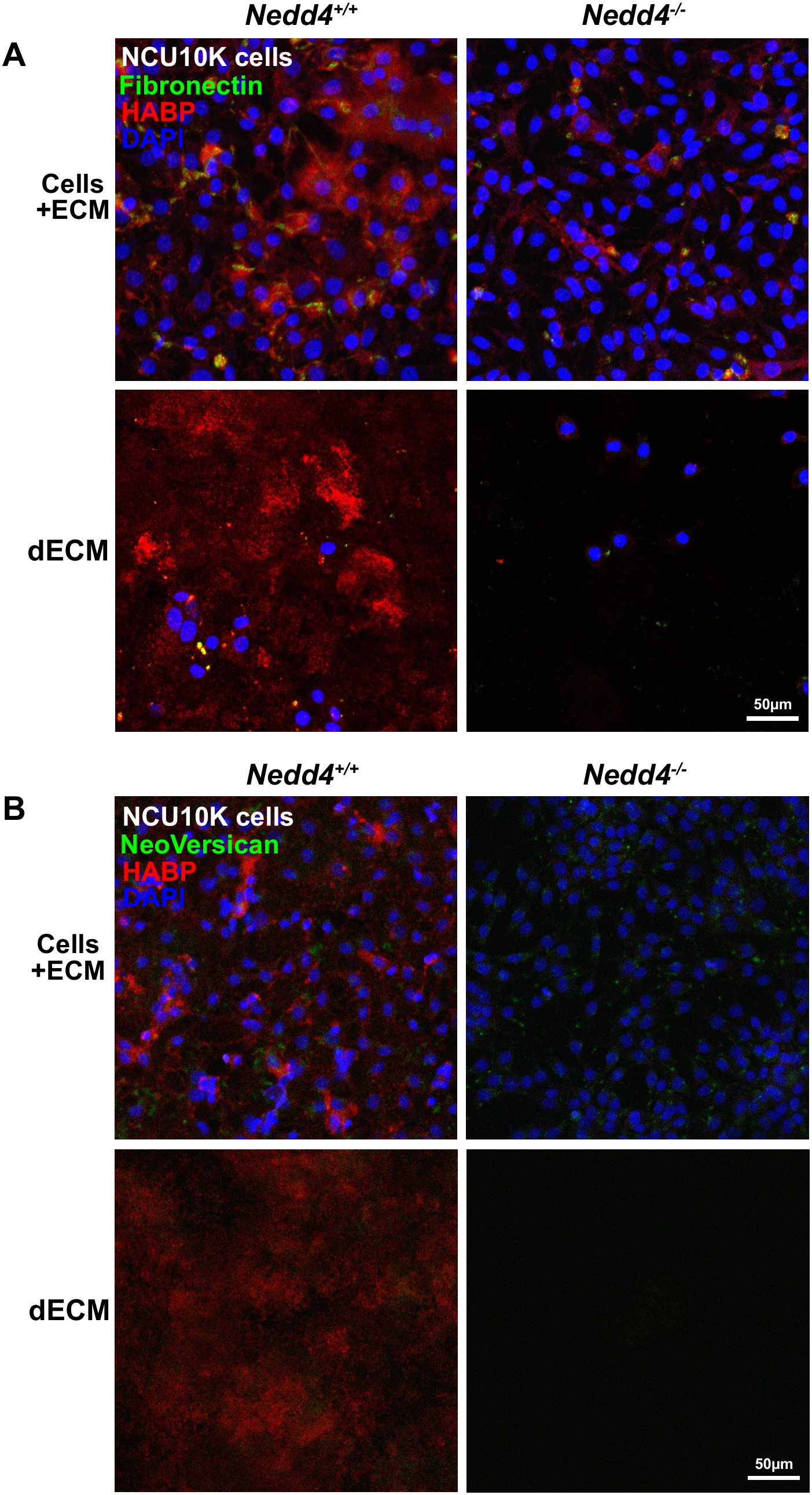
NCU10K cells produce negligible Fibronectin and NeoVersican. **A:** Immunostaining of NCU10K *Nedd4^+/+^* or *Nedd4^-/-^* fixed cells or decellularised ECM (dECM) for Fibronectin and HABP. Minimal Fibronectin is evident, irrespective of genotype or in whole cells versus dECM. **B:** Immunostaining of NCU10K *Nedd4^+/+^* or *Nedd4^-/-^* fixed cells or decellularised ECM (dECM) for NeoVersican and HABP. Minimal NeoVersican is evident, irrespective of genotype or in whole cells versus dECM.

**Supplementary Figure 7:**
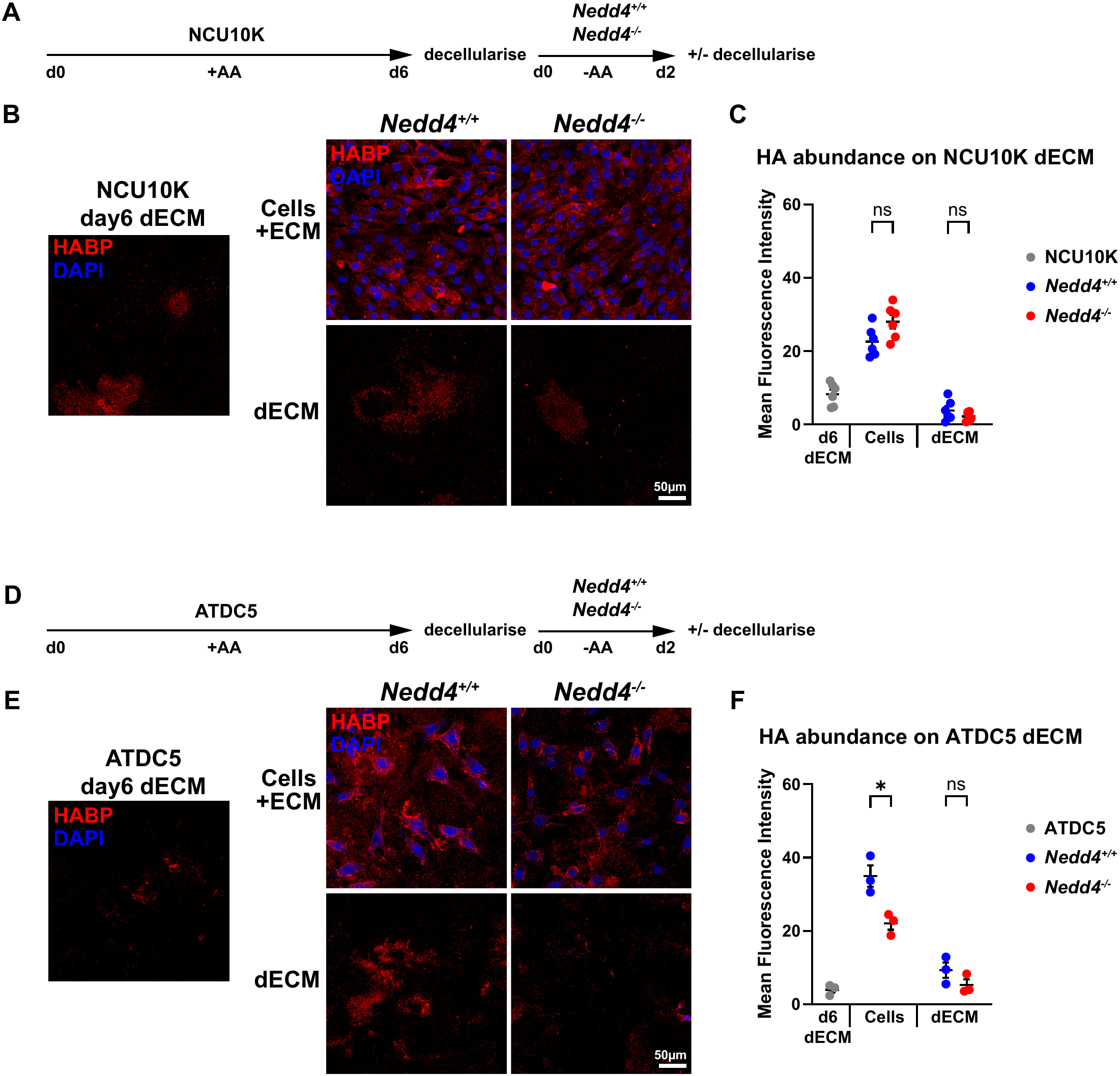
Using cell-derived ECM from NCU10K and ATDC5 cells to assess modulation of extracellular HA by *Nedd4^+/+^* and *Nedd4^-/-^* cells. **A:** Diagram of experimental procedure, culture conditions and timing of cell fixation/decellularisation. NCU10K parental cells were cultured for 6 days with ascorbic acid then decellularised to generate dECM. *Nedd4^+/+^* or *Nedd4^-/-^* cells were then plated onto the dECM and cultured for 2 days without ascorbic acid, then either fixed intact or decellularised. **B:** Staining of fixed cells or dECM with biotinylated hyaluronic acid binding protein (HABP) to label areas of HA. Staining of NCU10K-derived dECM only after 6 days prior to plating of cells on top, or following culture with *Nedd4^+/+^* or *Nedd4^-/-^* cells for 2 days, as fixed cells or dECM. **C:** Graph of HA mean fluorescence intensity (MFI), quantified from images as in (B). ns, not significant; 2 way ANOVA multiple comparisons. **D:** Experimental procedure for generation of dECM substrate from ATDC5 cells, followed by culture with *Nedd4^+/+^* or *Nedd4^-/-^* cells. **E:** HABP staining of ATDC5-derived dECM only after 6 days prior to plating of cells on top, or following culture with *Nedd4^+/+^* or *Nedd4^-/-^* cells for 2 days, as fixed cells or dECM. **F:** Graph of HA MFI, quantified from images as in (E). * p<0.05; ns, not significant; 2 way ANOVA multiple comparisons.

